# DNA methylation marks associated with body composition in children from India and the Gambia - findings from the EMPHASIS study

**DOI:** 10.1101/2025.05.15.654252

**Authors:** Prachand Issarapu, Manisha Arumalla, Elie Antoun, Chiara di Gravio, Kate Ward, Caroline H D Fall, Andrew M Prentice, Giriraj Ratan Chandak, Matt J Silver, the EMPHASIS study group

**Affiliations:** Genomic Research on Complex Diseases (GRC Group), CSIR-Centre for Cellular and Molecular Biology, Hyderabad, India; MRC Unit The Gambia at the London School of Hygiene and Tropical Medicine, The Gambia; London School of Hygiene and Tropical Medicine, London, UK; School of Medicine, University of Southampton, Southampton, UK; MRC Lifecourse Epidemiology Centre, Human Development and Health, University of Southampton, Southampton, UK

**Keywords:** Epigenetics, DNA methylation, Body composition EMPHASIS study, Epigenome Wide Association Study (EWAS)

## Abstract

**Background:** Differences in body composition during childhood can influence long-term health, with notable links to cardiometabolic disorders in later life. While genetic associations with body composition traits are well-studied, less is known about the role of epigenetic mechanisms, particularly in low- and middle-income countries where the burden of cardiometabolic disease is high. We investigated links between DNA methylation and three compartments of body composition: fat mass, lean mass, and bone measures using data from children enrolled in the *Epigenetic Mechanisms linking Pre-conceptional nutrition and Health Assessed in India and Sub-Saharan Africa* (EMPHASIS) study.

**Results:** We conducted an epigenome-wide association study (EWAS) of 11 body composition traits assessed through dual-energy X-ray absorptiometry in children from India (age = 5-7 years, n = 686) and The Gambia (age = 7-9 years, n = 289), with blood DNA methylation measured at approximately 800,000 CpGs sites on on the Illumina EPIC array. Cohort-specific analysis identified 15 unique differentially methylated CpGs (dmCpGs) associated with traits across all three compartments of body composition (p<3.6x10^-8^). Cross-cohort meta-analysis revealed 4 loci associated with lean mass and bone area. Notably, dmCpGs mapping to the *SOCS3* gene, previously linked to height in Indian, African and European populations, were associated with lean mass and bone area in both the Indian cohort and combined meta-analyses. Region-level EWAS identified differentially methylated regions (DMRs), linked to lean mass and bone area mapping to *SOCS3* and *P4HB* genes overlapping identified dmCpGs associated with the same phenotypes. Other DMRs mapped to genes including *BPNT1, RNU5F-1, LTA, HIF1A, HIF1A-AS1, MTHFD1*, and *TRIM72* were associated with multiple traits.

**Conclusion:** We report novel DNA methylation signatures associated with body composition traits in children from two low- and middle-income countries, highlighting a potential role for epigenetic mechanisms in shaping early-life body composition.

## Background

Body composition in childhood plays an important role in shaping long-term health outcomes^1,2^. The three-compartment model, which evaluates fat mass (FM), fat-free mass or lean mass (LM), and bone mineral content (BMC), is used to estimate the body’s total functional capacity and overall health status^3^. Deviations in these measures can significantly increase the risk of various health conditions both in childhood and later in life^1,4^. For example, excessive fat mass and insufficient lean mass during childhood are strongly linked to developing type 2 diabetes (T2D), hypertension and cardiovascular disease in adulthood^5–8^, and obesity in childhood heightens the risk of several cancers, orthopaedic complications and cardiometabolic disorders^9,10^.

The prevalence of body composition-related disorders has been rising worldwide^11,12^, with large differences between high-income countries (HICs) and low- and middle-income countries (LMICs)^11^. Furthermore, LMICs such as India and The Gambia face a dual burden of malnutrition, with undernutrition coexisting with rising incidences of overweight and obesity. In the United States, 19.3% of children and adolescents aged 2-19 years were classified as obese in 2017-2018^13^. In contrast, the prevalence of childhood obesity in India is estimated to be between 10-30%, varying by region^14^, while in The Gambia, urban children show increasing trends of overweight and obesity^15^.

Genome-wide association studies (GWAS) have helped to elucidate molecular mechanisms underlying body composition traits,^16,17^ with genetic variants identified for key traits such as body mass index (BMI)^18^, adiposity^19^, and lean mass^20^. In addition to genetic influences, environmental factors including lifestyle, physical activity, and dietary practices play a significant role in shaping body composition^21^, and there is growing interest in the potential for epigenetic mechanisms to mediate genetic and environmental effects on body composition and related traits^22–24^. For example, epigenome-wide association studies (EWAS) have identified associations between DNA methylation (DNAm) and adiposity^25^, BMI^26^, bone growth^27^, obesity^28^ and diabetes^29^. However, there is limited data from LMICs, where environmental and nutritional factors may have a greater impact on body composition^30^.

The *Epigenetic Mechanisms linking Periconceptional Nutrition and Health Assessed in India and sub-Saharan Africa* (EMPHASIS) study^31^ combines deep phenotyping with genetic and DNAm profiling in children from two LMICs, India and The Gambia. In our previous analyses, we reported EWAS findings linking DNAm with periconceptional maternal nutrition^32^, cardiometabolic risk markers^33^, and childhood height^30^. Here, we present findings from an EWAS investigating links between DNAm and body composition.

## Results

Baseline characteristics for the Indian and Gambian cohorts analysed in this study are presented in Table 1. Further details are provided in Methods.

**Table 1:** Baseline characteristics for EMPHASIS cohorts analysed in this study.

| **Child Parameters** | **Indian (n=686)** |  | **Gambian (n=284)** |  |
| --- | --- | --- | --- | --- |
| Sex (F) | 309 (45%) |  | 155 (55%) |  |
| Age (yrs) | 5.75 | (5.63,5.98) | 9.01 | (8.63,9.22) |
| Height (cm) | 109.4 | (106.40,113.00) | 130.40 | (125.70,133.60) |
| BMI (kg/m<sup>2</sup>) * | 13.28 | (12.62,13.80) | 14.24 | (13.52,15.13) |
| Lean mass (kg) | 12.65 | (11.58,13.68) | 18.34 | (16.56,19.93) |
| Fat mass (kg) | 2.24 | (1.77,2.92) | 4.69 | (4.02,5.51) |
| Android fat (kg) | 0.15 | (0.12,0.20) | 0.12 | (0.10,0.17) |
| Gynoid fat (kg) | 0.60 | (0.49,0.74) | 0.66 | (0.48,0.83) |
| Lean mass index (kg/m<sup>2</sup>)* | 10.57 | (10.04,11.10) | 10.89 | (10.17,11.55) |
| Fat mass index (kg/m<sup>2</sup>)* | 1.88 | (1.48,2.45) | 2.77 | (2.41,3.25) |
| Body fat percentage (%)* | 14.41 | (11.62,17.91) | 19.96 | (17.41,22.56) |
| Bone area (cm<sup>2</sup>) | 528.5 | (469.11,582.90) | 1107.32 | (1025.30,1165.90) |
| Bone mineral content (g) | 304.10 | (261.80,348.80) | 665.00 | (586.30,744.60) |
| Bone mineral density (g/cm<sup>2</sup>)* | 0.58 | (0.55,0.60) | 0.60 | (0.56, 0.64) |
| % stunted (n) | 686 | 15.3% (105) | 289 | 7.6% (22) |
| % wasted (n) | 686 | 37.2% (255) | 289 | 22.5% (65) |
| **Maternal Parameters** |  |  |  |  |
| Age at recruitment (yrs) | 25.00 | (22.00,27.00) | 30.04 | (25.87,34.61) |
| BMI (kg/m<sup>2</sup>) | 19.56 | (17.70,22.46) | 20.79 | (19.34,22.98) |
| Nutritional intervention<br>(supplementation/control) | 321/365 |  | 135/149 |  |

Median values are presented with IQR values in parentheses. n: number of participants included in the analysis; F: female; BMI: Body Mass Index; \*: Derived measures.

We conducted EWAS in each cohort to identify associations between peripheral blood DNAm measured on the Illumina EPIC array and 11 phenotypes covering the three compartments of body composition: lean mass (2), fat mass (5 measures) and bone measures (3) by dual-energy X-ray absorptiometry, together with BMI (Figure 1; Methods). All outcomes were analysed as residuals adjusted for age and sex (Methods). Differentially methylated CpGs (dmCpGs) were identified in each cohort separately using robust linear regression models implemented in the *ewaff* package in R. This was followed by meta-analysis of both cohorts combined using *METAL*^*34*^. Significant associations were defined as those passing a methylome-wide threshold p < 3.6x10^-8^, accounting for multiple testing^35^, applied independently to each outcome. Regional methylation analysis to identify differentially methylated regions (DMRs) was conducted on each cohort separately using *dmrff*^*36*^ in R. Significant associations were defined as those passing a Bonferroni-adjusted significance threshold p < 0.05, applied independently to each outcome. Further details on all analysis steps are given in Methods.

**Figure 1:**
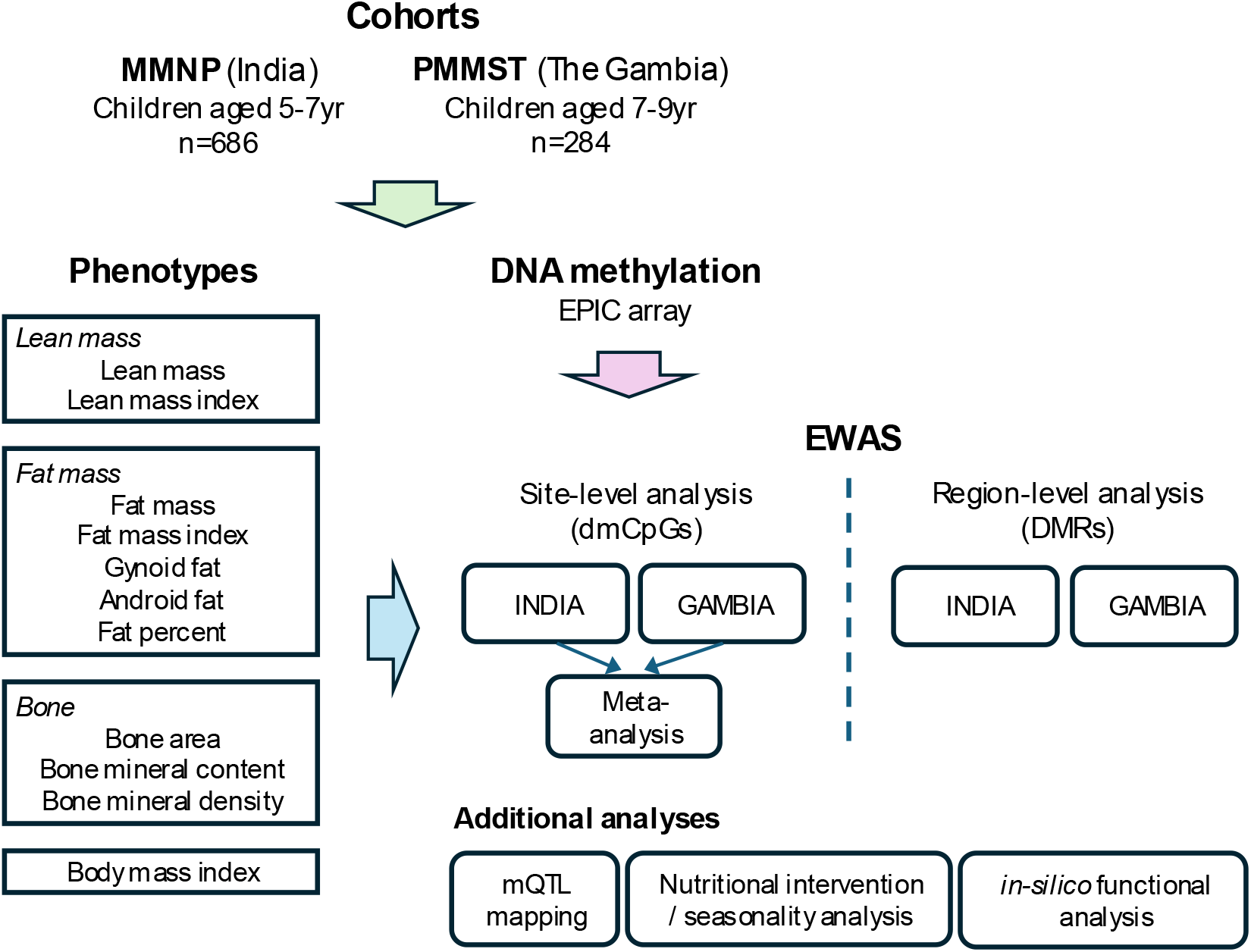
Schematic representation of study design. Body composition and body mass index phenotypes, together with genome-wide DNA methylation values were obtained for children from Indian and Gambian cohorts in the EMPHASIS study. Site-level and Region-level associations were identified through epigenome-wide analysis. Additional analyses investigated genetic influences and *in-silico* functional analysis. EMPHASIS: Epigenetic Mechanisms linking Periconceptional Nutrition and Health Assessed in India and sub-Saharan Africa; MMNP: Mumbai Maternal Nutrition Project; PMMST: Periconceptional Maternal Micronutrient Supplementation Trial; EWAS: Epigenome-wide association study; dmCpGs: differentially methylated CpGs; DMRs: differentially methylated regions; mQTLs: methylation quantitative trait loci.

### Indian EWAS

Significant dmCpGs in the India-only EWAS are presented in the top half of Table 2 and Figure 2A.

**Table 2:**
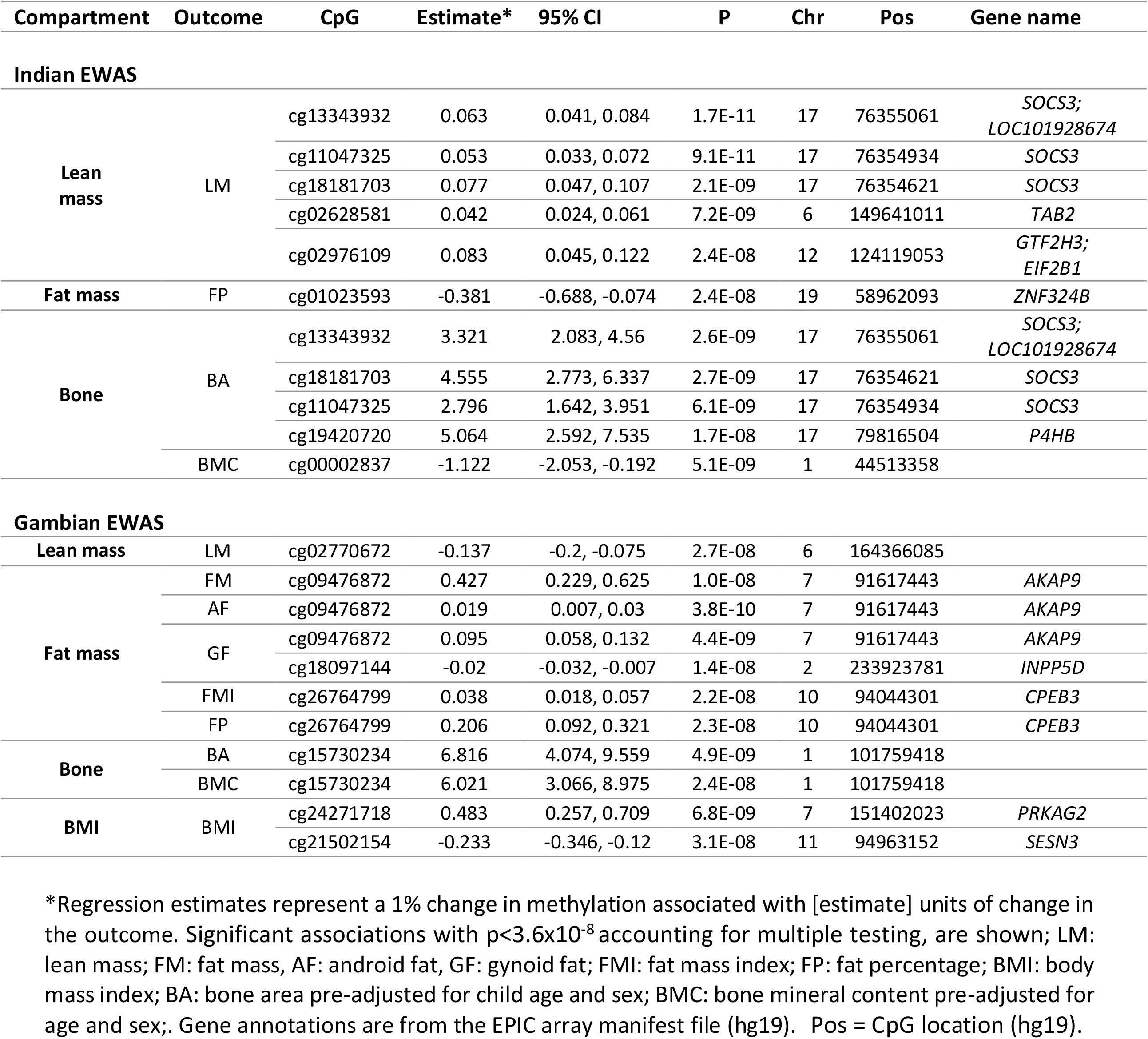
dmCpGs associated with body composition in Indian and Gambian children.

**Figure 2:**
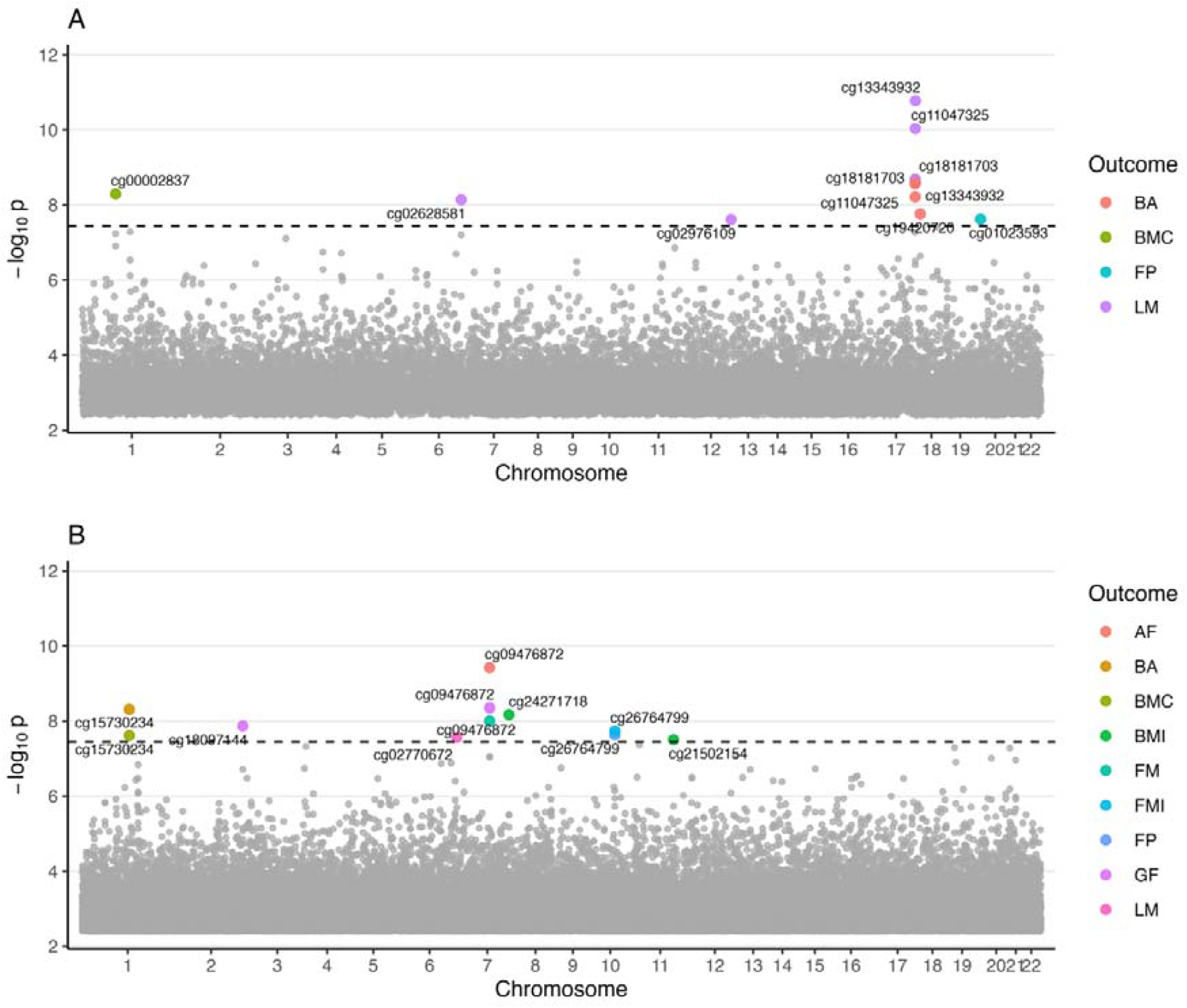
Identification of dmCpGs in Indian **(A)** and Gambian **(B)** cohorts. Significant CpGs are coloured according to outcome as indicated. The horizontal dashed line represents the p-value threshold for significant methylome-wide associations accounting for multiple testing. BA = bone area; BMC = bone mineral content; BMI = body mass index; FP = fat percent; LM = lean mass; FMI = fat mass index; AF = android fat; GF = gynoid fat.

#### Lean mass

Five dmCpGs were associated with lean mass. The top three were located within the suppressor of cytokine signalling 3 (*SOCS3*) gene: a 1% change in methylation was associated with a 0.06kg in increase in lean mass at at cg13343932 (95% confidence interval: 0.041, 0.084; p = 1.7x10^-11^), and with 0.05kg (0.033, 0.072, p=9.1x10^-11^) and 0.077kg (0.047, 0.11, p=2.1x10^-9^) increases in lean mass for cg11047325 and cg18181703 respectively. cg02628581, annotated to the TGF-Beta Activated Kinase 1 (*MAP3K7*) Binding Protein 2 (*TAB2*) gene, was associated with a 0.04 kg increase in lean mass per 1% change in methylation (95% CI: 0.02, 0.06; p = 7.2x10^-09^). cg02976109 (95% CI: 0.05, 0.12, p = 2.4x10^-8^) overlaps with the first intron of General Transcription Factor IIH Subunit 3 (*GTF2H3*) gene and the 5’UTR of Eukaryotic Translation Initiation Factor 2B Subunit Alpha (*EIF2B1)* and was associated with an increase of 0.08 kg in lean mass per 1% change in methylation. No significant associations with lean mass index were observed.

#### Fat mass

There were no significant associations with fat mass, fat mass index or android and gynoid fat measures.cg01023593 within the Zinc Finger Protein 324B (*ZNF324B*) gene was associated with fat percent, with a decrease of 0.38 units per 1% increase in methylation (95% CI: -0.69, -0.07; p = 2.4x10^-08^).

#### Bone measures

Four dmCpGs were associated with bone area. Three of these were located within the *SOCS3* gene: cg13343932 (3.32 cm^2^ per 1% increase in methylation; 95% CI: 2.08, 4.56; p = 2.6x10^-09^), cg18181703 (4.56 cm^2^; 95% CI: 2.77, 6.34; p = 2.7x10^-09^), and cg11047325 (2.80 cm^2^; 95% CI: 1.64, 3.95; p = 6.1x10^-09^). The fourth dmCpG in Prolyl 4-Hydroxylase Subunit Beta (*P4HB*), cg19420720 was associated with an increase of 5.06 cm^2^ (95% CI: 2.59, 7.54; p = 1.7x10^-08^). cg00002837 was associated with a decrease of 1.12 g in bone mineral content per 1% increase in methylation (95% CI: -2.05, -0.19; p = 5.1x10^-09^). No significant associations with bone mineral density were observed.

### Gambian EWAS

Significant dmCpGs in the India-only EWAS are presented in the bottom half of Table 2 and Figure 2B.

#### Lean mass

One dmCpG, cg02770672, mapping to an intergenic region was associated with lean mass in this cohort, with a 1% increase in methylation corresponding to a 0.14 kg decrease in lean mass (95% CI: - 0.21, -0.07; p = 2.7x10^-08^). No significant associations with lean mass index were observed.

#### Fat mass

A 1% increase in methylation at cg09476872, mapping to the A-Kinase Anchoring Protein 9 (*AKAP9*) gene was associated with a 0.43 kg increase in fat mass (95% CI: 0.23, 0.63; p = 1.0x10^-08^). This CpG was also associated with a 0.02 kg increase in android fat (95% CI: 0.01, 0.03]; p = 3.8x10^-10^), and a 0.10 kg increase in gynoid fat (95% CI: 0.60, 0.13; p = 4.4x10^-09^). cg18097144, mapping to the Inositol Polyphosphate-5-Phosphatase D (*INPP5D*) gene, was also associated with GF, with a 1% increase in methylation corresponding to a decrease of 0.02 kg (95% CI: -0.03, -0.01; p = 1.4x10^-08^).

cg26764799, within the Cytoplasmic Polyadenylation Element Binding Protein 3 (*CPEB3*) gene was associated with both FMI and FP, with estimated increases of 0.04 kg/m^2^ (95% CI: 0.02, 0.06; p = 2.2x10^-08^) and 0.21 percent (95% CI: 0.09, 0.32; p = 2.3x10^-08^) per 1% increase in methylation respectively.

#### Bone measures

cg15730234 was associated with an increase of 6.81 cm^2^ (95% CI: 4.07, 9.56; p = 4.9x10^-09^) in bone area, and with an increase of 6.02 g (95% CI: 3.07, 8.98; p = 2.4x10^-08^) in bone mineral content. No significant associations with bone mineral density were observed.

#### Body Mass Index

Two CpGs showed significant associations with BMI. A 1% increase in methylation at cg24271718 in the Protein Kinase AMP-Activated Non-Catalytic Subunit Gamma 2 (*PRKAG2*) gene was associated with an increase of 0.48 kg/m^2^ in BMI (95% CI: 0.26, 0.71; p = 6.8x10^-09^). The same increase in methylation a cg21502154 in the Sestrin 3 (*SESN3*) gene was associated with a reduction of 0.213 kg/m^2^ in BMI (95% CI: -0.35, -0.12; p = 3.1x10^-08^).

Note no dmCpGs overlapped both cohorts.

### Meta analysis

Meta-analysis of summary statistics from individual cohort EWAS identified significant associations with lean mass and bone area adjusted for age and sex (Table 3, Figure 3).

**Table 3:** Combined meta-analysis of cohort-level CpG associations.

| Outcome | Name | Effect | StdErr | P | Direction | HetISq | HetPVal | Chr | Pos | Gene Name |
| --- | --- | --- | --- | --- | --- | --- | --- | --- | --- | --- |
| Lean mass | cg13343932 | 0.066 | 0.010 | 7.7E-12 | ++ | 19.9 | 0.264 | 17 | 76355061 | *SOCS3*;<br>*LOC101928674* |
|  | cg11047325 | 0.055 | 0.009 | 7.3E-11 | ++ | 41 | 0.193 | 17 | 76354934 | *SOCS3* |
|  | cg18181703 | 0.079 | 0.014 | 3.3E-09 | ++ | 60.4 | 0.112 | 17 | 76354621 | *SOCS3* |
|  | cg25345365 | 0.045 | 0.013 | 7.0E-09 | ++ | 0 | 0.987 | 11 | 114050114 | *ZBTB16* |
| Bone area | cg25345365 | 2.939 | 0.718 | 1.8E-09 | ++ | 0.2 | 0.317 | 11 | 114050114 | *ZBTB16* |

Cohort-level EWAS summary statistics from Indian and Gambian cohort were meta-analysed using an inverse variance fixed-effects model. Significant associations with $p < 3.6 \times 10^{-8}$ accounting for multiple testing, are shown. HetISq and HetPVal values estimate the heterogeneity in effect sizes between cohorts. Gene names are from the EPIC array manifest file. Pos = CpG location (hg19). StdErr = standard error. Direction = direct of effect in each cohort.

**Figure 3:**
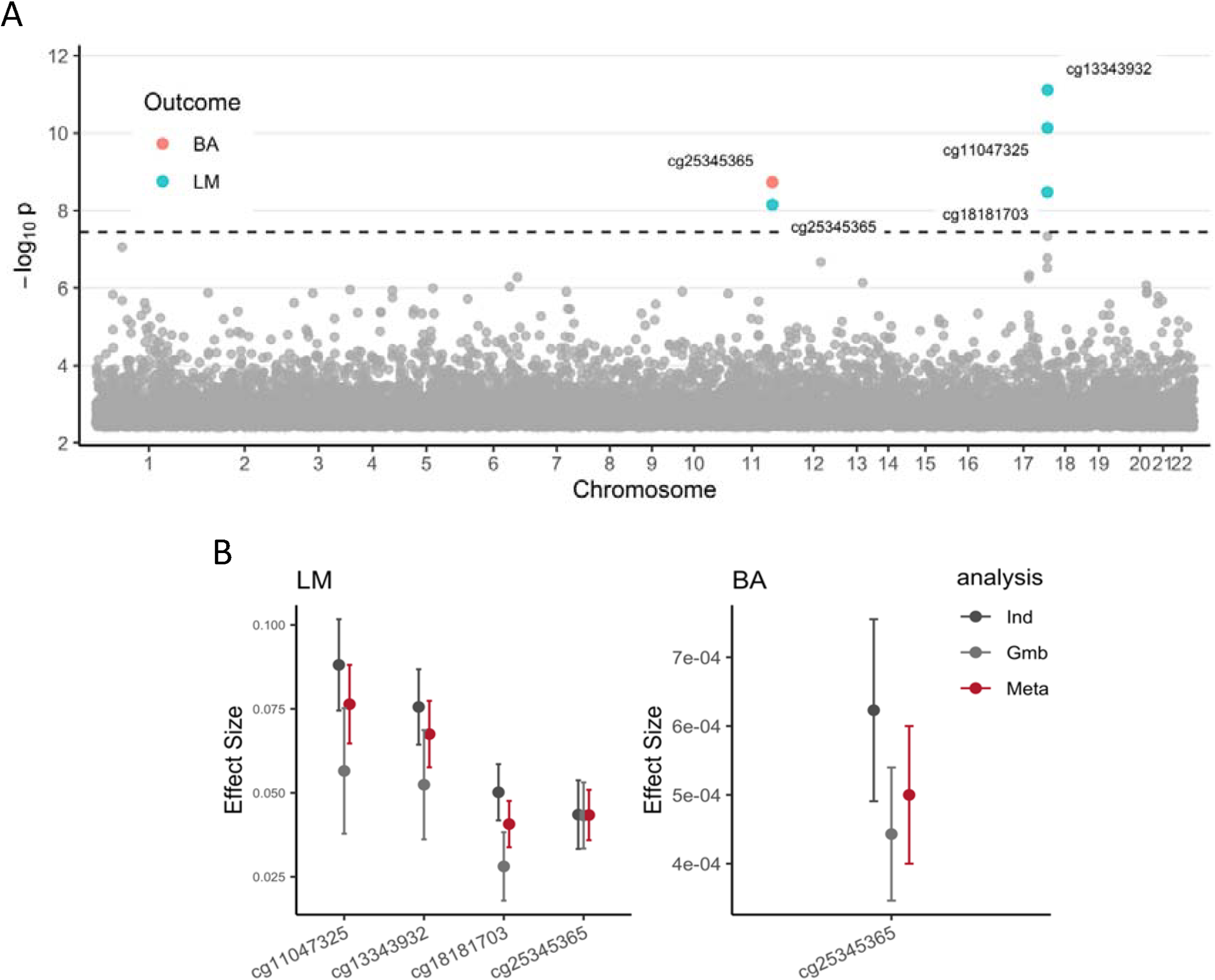
Meta-analysis of summary statistics from Indian and Gambian EWAS. **(A)** Manhattan plot showing significant dmCpGs. These are coloured according to their corresponding outcomes. **(B)** Forest plots showing effect sizes for significant CpGs in the individual cohort and meta-analysis. Error bars represent standard errors. LM: Lean mass; BA: Bone area; Ind: India; Gmb: Gambia.

Four dmCpGs were associated with lean mass, including three CpGs identified in the Indian cohort dmCpG analysis mapping to the *SOCS3* gene. Heterogeneity scores^37^ ranging from 20 to 60 suggested that the meta-analysed associations may be driven by the strong association in the Indian cohort. In contrast, a fourth dmCpG, cg25345365, within the zinc finger and BTB domain containing 16 (*ZBTB16*) gene was not identified in separate cohort analyses and showed a consistent direction of effect across both cohorts in the meta-analysis, with no evidence of heterogeneity. This CpG was also associated with bone area in the meta-analysis, again with no evidence of heterogeneity.

### Sensitivity analyses

#### Maternal micronutrient supplementation and Gambian season of conception

Mothers in both participating cohorts received pre- and peri-conceptional micronutrient supplementation^38^. In a previous report in the EMPHASIS study, we noted an effect of intervention on offspring DNA methylation at several loci^32^. We found no effect of the interventions on body composition-associated dmCpGs in either cohort (Supplementary Table 1). We additionally tested for an effect of season of conception in the Gambian cohort^32^ and found no significant association (Supplementary Table ).

### Differentially methylated regions

We explored region-level DNAm associations for all 11 body composition measures, separately in Indian and Gambian cohorts.

We identified 30 DMRs associated with multiple body composition outcomes in the Indian cohort (Supplementary Table 3). Notably, lean mass, bone area and bone mineral content phenotypes shared associations with DMRs mapping to common genes including Tripartite Motif Containing 72 (*TRIM72*), Cyclin Dependent Kinase Like 1 (*CDKL1*) and Lymphotoxin Alpha (*LTA*), while a region mapping to Zinc Finger Protein 772 (*ZNF772*) was associated with lean mass and fat percent phenotypes. Several trait-specific DMRs were also identified. DMRs mapping to *SOCS3* and *P4HB* genes overlapped CpGs identified in the Indian dmCpG analysis.

We identified 22 DMRs in the Gambian cohort (Supplementary Table 4). Notably, DMRs mapping to Bisphosphate 3’-Nucleotidase 1 (*BPNT1*) and RNA, U5F Small Nuclear 1 (*RNU5F-1*) were associated with fat mass, fat mass index, fat percent and body mass index, and a DMR located in Methylenetetrahydrofolate Dehydrogenase/ MicroRNA 548az (*MTHFD1/MIR548AZ*), was associated with BMI, bone area and android fat. Additionally, the *HIF1A*/*HIF1A-AS1* locus was associated with android fat and BMI and there were multiple trait-specific DMRs identified in this cohort.

There were no overlapping DMRs between Indian and Gambian cohorts.

### Influence of genetic variants on body composition-associated dmCpGs

We investigated the potential influence of genetic variation on dmCpGs by conducting a methylation quantitative trait locus (mQTL) analysis on SNPs ±1Mb from the dmCpGs. This analysis was performed after SNP pruning to identify SNPs tagging linkage disequilibrium blocks. We found no significant associations with dmCpGs in either cohort after adjustment for multiple testing.

### Functional annotations

We explored the potential functional relevance of dmCpGs identified in this study by checking for their associations with disease outcomes in the catalogue of previously reported EWAS results. Notably dmCpGs mapping to *SOCS3* (cg18181703, cg13343932, cg11047325) have previously been linked to multiple traits including obesity and rheumatoid arthritis. cg19420720 (mapping to *P4HB*) and cg00002837 (mapped to intergenic region) have been linked to pre-eclampsia and c-reactive protein levels respectively (Supplementary Table 5). We further investigated the involvement of identified body composition associated genes in the known molecular pathways using Gene Ontology (GO) and KEGG pathway analyses. Of the 43 unique genes identified, 38 could be mapped to Entrez IDs. However, no significant enrichment for GO terms or KEGG pathways were observed.

## Discussion

We investigated links between DNAm and body composition traits in children from two LMIC cohorts in the EMPHASIS study. Our study is notable for its comprehensive analysis of genome-wide DNAm signatures associated with four key DEXA measures: fat mass, lean mass, bone density and bone mineral content. Through a combination of cohort-specific and combined cohort meta-analysis, we identified 17 unique CpGs, some of which were trait-specific while others overlapped across multiple traits. We also identified 52 DMRs. Our study provides potential insights into molecular mechanisms underpinning body composition traits in these understudied populations.

Methylation at multiple dmCpGs in the *SOCS3* gene were associated with lean mass (Indian and combined cohort meta-analysis) and bone area (Indian cohort analysis only). *SOCS3* is a known regulator of both inflammatory and growth-related pathways and may play a central role in coordinating muscle and bone development^39^. It modulates the JAK/STAT signalling pathway, regulates osteoblast and osteoclast activity critical for bone remodelling^39 40^, and influences cytokine-driven corticalization, which contributes to skeletal integrity ^41^. In muscle, *SOCS3* is upregulated in response to inflammation and elevated IL-6 levels, where it inhibits insulin/IGF-1/Akt signalling, suppresses protein synthesis, and promotes muscle atrophy through increased degradation^42,43^. These pleiotropic roles suggest that *SOCS3* methylation could be a key epigenetic regulator linking immune signalling to structural body composition during childhood.

We previously observed a strong association of *SOCS3* methylation with height in these cohorts, with evidence of a causal effect in European populations^30^. This is consistent with findings from this study, given the strong correlation between height and body composition measures^44^, and implies that *SOCS3m* may exert its primary influence on overall growth, which in turn is reflected in bone and muscle measures.

*ZBTB16* identified in the meta-analysis plays a role in body composition and related traits through its influence on metabolic regulation, energy expenditure, and thermogenesis^45^. Induced by cold exposure, ZBTB16 promotes energy expenditure and affects adipogenesis, cardiac hypertrophy, lipid levels and insulin sensitivity^46^.

Our analysis of DMRs in the Indian cohort identified several other genes with evidence of links to body composition traits and related metabolic outcomes. TRIM72 is involved in muscle repair and regeneration and helps suppress chronic muscle inflammation, a contributor to insulin resistance and obesity^47^. LTA regulates immune responses and has been implicated in chronic inflammatory diseases that affect fat distribution and muscle health^48,49^. TAB2 acts as a central mediator of inflammation by regulating NF-κB and MAPK signalling pathways, both essential for metabolic homeostasis and growth^50,51^. *SH2B3* negatively regulates cytokine signalling pathways and adipose tissue inflammation^52^ and has been linked to inflammation-driven hypertension^53^. P4HB, involved in autophagy-mediated pathways, is associated with proinsulin maturation and diabetic nephropathy, highlighting its role in inflammation and metabolic regulation^54,55^. Collectively, these genes point to a shared regulatory axis linking inflammation, growth, and body composition, underscoring their potential relevance to early-life metabolic programming.

DMR analysis in the Gambian cohort also identified genes with potential links to body composition traits and metabolic health. Notably, *INPP5D* and *PRKAG2* have been previously linked to cellular signalling and metabolic regulation. *INPP5D* identified with gynoid fat plays a role in regulating immune responses and lipid metabolism^56,57^, and has also been linked to Alzheimer’s disease^56,58^. *PRKAG2* encodes a regulatory subunit of AMP-activated protein kinase (*AMPK*), a master regulator of cellular energy balance. It influences glucose and lipid metabolism and has been associated with cardiac hypertrophy, obesity, and T2D^59–61^. *HIF1A*, associated with android fat and BMI in our study, regulates cellular adaptation to low oxygen levels, and controls genes involved in energy metabolism, angiogenesis, and erythropoiesis^62^. Its role in metabolic processes may impact body composition, particularly under hypoxic conditions that influence fat and muscle tissue responses^63^, and variants in *HIF1A* have been linked to preeclampsia^64^, diabetes^65^, cardiovascular diseases and cancers^63,66^. *SLC16A3* encodes a monocarboxylate transporter involved in lactate transport, essential for muscle metabolism and energy production^67,68^ and plays a significant role in metabolic conditions such as obesity and diabetes, where lactate clearance is compromised^67,69^. *MTHFD1*, involved in folate metabolism and nucleotide biosynthesis, plays a role in cell growth and energy metabolism^70^ and mutations in this gene have been linked to megaloblastic anaemia^70^. Together, these findings suggest that epigenetic regulation of several genes related to energy balance, glucose metabolism and immune signalling may be linked to body composition traits and associated metabolic disease.

Separate analyses of Indian and Gambian cohorts revealed distinct epigenetic associations with no overlapping dmCpGs or DMRs. In the Indian cohort, many associated genes have been linked to inflammation response, while in the Gambian cohort, several were involved in hypoxia response and energy balance. This underscores the importance of conducting EWAS in diverse ancestries as population-specific epigenetic signatures could arise in response to unique environmental exposures in populations with varying genetic backgrounds.

This study has several limitations. Our analysis of blood samples for DNAm analysis means that we were unable to analyse methylation patterns in other tissues, such as adipose or muscle, which may be more relevant to body composition traits. Additionally, the Indian cohort analysis was better powered with a larger sample size which could in part explain inter-cohort differences, and the overlap between Indian cohort and meta-analysis results. Furthermore, the associations that we report are correlative, so that we cannot infer causality or account for effects from unmeasured confounders including genetic variants. Our findings require replication in independent cohorts and further functional validation to clarify their biological relevance. Finally, our analysis was restricted to the small fraction of the methylome captured by EPIC arrays, potentially missing important signals outside the assayed regions.

## Conclusions

Our study provides insights into the epigenetic regulation of body composition in children from two LMIC populations with an analysis of genome-wide DNA methylation signatures associated with fat mass, lean mass, and bone outcomes. We identify differentially methylated CpGs and regions across multiple outcomes, the majority of which are cohort-specific, highlighting the potential role of genetic heterogeneity and differential responses to distinct environmental exposures between the two populations. These findings highlight methylation loci of potential relevance to body composition traits that warrant further investigation in the context of growth and metabolic health in diverse populations.

## Methods

### Study cohorts

The cohorts included in this study are part of the EMPHASIS study (ISRCTN14266771), which is a follow-up of children born to mothers who participated in two independent randomized controlled trials of peri-conceptional maternal micronutrient supplementation in two undernourished communities. The Mumbai Maternal Nutritional Project (MMNP: ISRCTN62811278)^38^ was conducted from 2006 to 2012 in the slums of Mumbai, India. The Peri-conceptional Maternal Micronutrients Supplementation Trial (PMMST: ISRCTN13687662)^38^ was conducted from 2006 to 2008 in West Kiang, The Gambia, in West Africa. In the Indian cohort, children were followed up at the age of 5-7 years, and the study included the first 709 children from the per-protocol group. In the Gambian cohort, children were followed up at 7-9 years of age, and all retraced children (n= 296) were included in the original study. Further details are provided in previous publications^38,71^.

### Body composition measures and pre-processing of phenotype data

Measures of lean mass, fat mass, and bone density and bone mineral content were obtained using dual-energy X-ray absorptiometry, using the Lunar Prodigy in India and Lunar iDXA in The Gambia. A total of 11 body size and composition measures were analysed in this study. The direct measures were lean mass, fat mass, gynoid and android fat mass, bone area and bone mineral density. The latter two bone measures were calculated for total body, excluding the head, as per international guidelines (https://iscd.org/learn/official-positions/pediatric-positions/). The derived measures were BMI (BMI = WT/HT^2^, where, WT and HT are child’s weight (in kilograms) and height (in metres), respectively); lean mass index (LMI = LM/HT^2^); fat mass index (FMI = FM/HT^2^); total body fat percent (body fat % = [FM/WT]*100); and bone mineral content (BMC = BA*BMD). All phenotype outcome variables were assessed for normality and log transformed wherever necessary (FM, FMI, FP, AF and GF) and outcome residuals were generated by regressing each outcome against age and sex.

### Pre-processing of methylation data

Detailed information on the data generation and pre-processing has been previously reported^72^. Briefly, genome-wide DNA methylation profiles were generated using the Illumina MethylationEPIC BeadChip (‘EPIC array’). Pre-processing of the methylation data was carried out using the *meffil* package from Bioconductor^73^. The data were normalized using the *functional normalization* method as part of the *meffil* pipeline^73^. Data from the Indian and Gambian cohorts were processed separately. Final analysed datasets comprised 686 samples with 803,210 probes for the Indian cohort and 289 samples with 802,283 probes for the Gambian cohort.

### Pre-processing of genotype data

Imputed genotype data were available for both the EMPHASIS study cohorts. Detailed information on the genotype data generation and imputation was reported previously^72^. Briefly, the array-derived data were pre-phased using SHAPEIT (version: v2.r900), and imputation was performed using IMPUTE2 (version 2.3.2) with the 1000 Genomes phase 3 reference panel. SNPs with a minor allele frequency (MAF) less than 10% and an IMPUTE ‘info’ metric less than 0.9 were excluded to ensure high imputation quality and reliability.

### Epigenome-wide association studies (EWAS)

#### Site-level differential methylation analysis

Epigenome-wide association analyses were carried out for each outcome using robust linear regression models implemented in the *ewaff* R package (https://github.com/perishky/ewaff) using default parameters. Log-transformed methylation beta values (M-values) were regressed against body composition outcome residuals, adjusting for child age, sex, and the first 10 surrogate variables (SVs) derived from the 200k most variable methylation probes. SVs were used to account for batch effects and variability in estimated blood cell composition while retaining the effect from the outcome variable of interest. Inflation of p-values was assessed through the genomic inflation factor (lambda; Supplementary tables 6 and 7) and visualized via quantile-quantile (QQ) plots (Supplementary Figures 1 and 2). Significant dmCpGs were defined as those passing a methylome-wide threshold p < 3.6x10^-8^, accounting for multiple testing^35^, applied independently to each outcome. For significant outcome-CpG associations, the models were re-run using methylation beta values (instead of M-values) to generate interpretable regression estimates where effect size is proportional to a unit change in the outcome. These are reported in the main text.

#### Region-level differential methylation analysis

To identify DMRs we used the R package *dmrff* (v1.0.0) (https://github.com/perishky/dmrff)^36^, which accounts for regional correlation between CpGs. A DMR was defined as the presence of two or more differentially methylated CpG sites within 1000 bp range.

### Meta-analysis

METAL^74^ (version: 2011-03-25; https://genome.sph.umich.edu/wiki/METAL) was used to meta-analyse individual EWAS summary statistics from each cohort. An inverse variance fixed effects model accounting for genomic inflation was implemented. Cross-cohort heterogeneity effects were assessed by the heterogeneity index (hetI^2^). We applied a Bonferroni significance threshold for identification of significant dmCpGs in the meta-analysis, as described above.

### Sensitivity analyses

We performed a sensitivity analysis on dmCpGs identified in single cohort and/or meta-analysis to check for effects of maternal supplementation carried out in the original Indian and Gambian studies and Gambian season of conception^35^. A Bonferroni corrected p-value threshold of 0.05 was used in each cohort.

### Influence of genetic variation on methylation signatures (mQTL analysis)

cis-mQTL analysis was conducted using the Gene-Environment-Methylation (*GEM*) package (v1.10.0) from R Bioconductor to assess SNP-methylation associations. An additive allelic dosage (0,1,2) regression model adjusted for child age, sex and the first 10 PCs derived from the 200K most variable methylation probes was used following the GEM pipeline. This analysis was carried out on unique dmCpGs identified in single cohort and/or meta-analyses and SNPs within ±1MB of the dmCpGs that were ‘pruned’ to account for linkage disequilibrium between the SNPs. The pruning was carried out in plink2 (https://www.cog-genomics.org/plink/2.0/) using --indep-pairwise with the following parameters: window size = 1000kb; step size (variant count) = 50; and r^2 = 0.2. This resulted in 971 and 3324 cis-SNPs in the Indian and the Gambian cohorts respectively. Significant cis-mQTLs were determined based on association p-values that passed a Bonferroni-adjusted significance threshold of p = 0.05/n_SNPs_tested/n_CpGs_tested; where ‘n_SNPs_tested’ represents the number of cis-SNPs and ‘n_CpGs_tested’ is the number of unique CpGs tested in the corresponding cohort.

### Functional annotation

To investigate the functional relevance of identified dmCpGs and DMRs, we conducted several *in silico* analyses using publicly available databases. First, we cross-referenced the identified dmCpGs against the EWAS Catalog (https://www.ewascatalog.org/) to see if they had been previously associated with body composition or related traits. Next, we used the *clusterProfiler* R package to identify enriched GO terms (BP, MG, CC) and KEGG pathways linked to genes containing these dmCpGs and DMRs, aiming to identify potential biological processes relevant to body composition.

## List of abbreviations

AF: Android fat
BA: Bone area
BMC: Bone mineral content
BMD: Bone mineral density
BMI: Body mass index
dmCpG: Differentially methylated CpG
DMR: Differentially methylated region
EWAS: Epigenome-wide association study
FDR: False discovery rate
FM: Fat mass
FMI: Fat mass index
GF: Gynoid fat
GO: Gene Ontology
KEGG: Kyoto Encyclopaedia of Genes and Genomes
LD: Linkage disequilibrium
LM: Lean mass
LMI: Lean mass index
LMIC: low-to-middle income countries
mQTL: Methylation quantitative trait loci

## Ethics approval and consent to participate

Ethics approval of the EMPHASIS study for the follow-up of the children in Mumbai (“SARAS KIDS”) was obtained from the Intersystem Biomedica Ethics Committee, Mumbai on 31 May 2013 (serial no. ISBEC/NR-54/KM/JVJ/2013). Ethics approval for the EMPHASIS study in The Gambia was obtained from the joint Gambia Government/MRC Unit The Gambia’s Ethics Committee on 19 October 2015 (serial no. SCC 1441). The EMPHASIS study is registered as ISRCTN14266771. Signed informed consent was obtained from parents and verbal assent from the children.

## Availability of data and materials

The data required for the results interpretation presented in this manuscript are provided within the manuscript and as supplementary material. Methylation and genotype data are available on request to the corresponding authors.

## Funding

The EMPHASIS study was funded under the Newton Fund initiative (MRC Grant No.: MR/N006208/1 and DBT Grant No.: BT/IN/DBT-MRC/DFID/24/GRC/2015–16) which was jointly funded by MRC, DFID and the Department of Biotechnology (DBT), Ministry of Science and Technology, India.

## Authors’ contributions

C.H.D.F., G.R.C., M.J.S. and A.M.P. conceived and designed the study. P.I., M.J.S., M.A., E.A., C.d.G., G.R.C., C.H.D.F., and A.M.P. devised the analysis strategy. P.I. performed bioinformatic and statistical analyses. P.I., G.R.C., and M.J.S. drafted the manuscript and all authors critically revised and approved the final manuscript.

## The EMPHASIS study group

**Lena Acolatse**, MRC Unit The Gambia at the London School of Hygiene and Tropical Medicine and University of Leeds, UK; **Meraj Ahmed**, CSIR-Centre for Cellular and Molecular Biology, Hyderabad, India; **Modupeh Betts**, MRC Unit The Gambia at the London School of Hygiene and Tropical Medicine and University of Liverpool, UK; **Giriraj R Chandak**, CSIR-Centre for Cellular and Molecular Biology, Hyderabad, India; **Harsha Chopra**, Centre for the Study of Social Change, Mumbai, India; **Cyrus Cooper**, MRC Lifecourse Epidemiology Centre, University of Southampton, UK; **Momodou K Darboe**, MRC Unit The Gambia at the London School of Hygiene and Tropical Medicine; **Chiara Di Gravio**, School of Medicine, University of Southampton, UK; **Caroline HD Fall**, MRC Lifecourse Epidemiology Centre, University of Southampton, UK; **Meera Gandhi**, Centre for the Study of Social Change, Mumbai, India; **Gail R Goldberg**, MRC Elsie Widdowson Laboratory, Cambridge, UK; **Prachand Issarapu**, CSIR-Centre for Cellular and Molecular Biology, Hyderabad, India and MRC Unit The Gambia at the London School of Hygiene and Tropical Medicine, UK; **Philip James**, London School of Hygiene and Tropical Medicine and MRC Unit The Gambia and London School of Hygiene and Tropical Medicine, UK; **Ramatoulie Janha**, MRC Unit The Gambia at the London School of Hygiene and Tropical Medicine; **Landing MA Jarjou**, MRC Unit The Gambia at the London School of Hygiene and Tropical Medicine; **Lovejeet Kaur**, CSIR-Centre for Cellular and Molecular Biology, Hyderabad, India; **Sarah H Kehoe**, MRC Lifecourse Epidemiology Centre, University of Southampton, UK; **Kalyanaraman Kumaran**, MRC Lifecourse Epidemiology Centre, University of Southampton, UK and CSI Holdsworth Memorial Hospital, Mysore, India; **Karen A Lillycrop**, University of Southampton, UK; **Mohammed Ngum**, MRC Unit The Gambia at the London School of Hygiene and Tropical Medicine; **Suraj S Nongmaithem**, CSIR-Centre for Cellular and Molecular Biology, Hyderabad, India; **Stephen Owens**, Newcastle University Trust Hospital, UK; **Ramesh D Potdar**, Centre for the Study of Social Change, Mumbai, India; **Andrew M Prentice**, MRC Unit The Gambia at the London School of Hygiene and Tropical Medicine and London School of Hygiene and Tropical Medicine, UK; **Ann Prentice**, MRC Unit The Gambia at the London School of Hygiene and Tropical Medicine, Elsie Widdowson Laboratory, Cambridge, UK and MRC Lifecourse Epidemiology Centre, University of Southampton; **Tallapragada Divya Sri Priyanka**, CSIR-Centre for Cellular and Molecular Biology, Hyderabad, India; **Ayden Saffari**, MRC Unit The Gambia at the London School of Hygiene and London School of Hygiene and Tropical Medicine, UK; **Sirazul Ameen Sahariah**, Centre for the Study of Social Change, Mumbai, India; **Sara Sajjadi**, CSIR-Centre for Cellular and Molecular Biology, Hyderabad, India; **Harshad Sane**, Centre for the Study of Social Change, Mumbai, India; **Smeeta Shrestha**, CSIR-Centre for Cellular and Molecular Biology, Hyderabad, India; **Matt J Silver**, MRC Unit The Gambia at the London School of Hygiene and Tropical Medicine and London School of Hygiene and Tropical Medicine, UK; **Ashutosh Singh Tomar**, CSIR-Centre for Cellular and Molecular Biology, Hyderabad, India; **Kate A Ward**, MRC Lifecourse Epidemiology Centre, University of Southampton, UK; **Dilip Kumar Yadav**, CSIR-Centre for Cellular and Molecular Biology, Hyderabad, India; **Chittaranjan S Yajnik**, Diabetes Unit, KEM Hospital and Research Centre, Pune, India.

## Acknowledgements

We thank the EMPHASIS study participants in Mumbai, India and West Kiang, The Gambia for their contribution to the original trials. We also thank laboratory personnel and field teams supported for this study in both countries. We are grateful to our steering committee members Caroline Relton, Partha P Majumder and Frank Dudbridge. We thank all the members of the EMPHASIS Study who have directly or indirectly contributed to the success of the EMPHASIS study.

## Consent for publication

Not applicable.

## Competing interests

The authors declare no competing interests.

## Authors’ information

Matt J Silver and Giriraj R Chandak: Joint corresponding authors.

## Notes

### Competing Interest Statement

The authors have declared no competing interest.

