## Supplementary_Figures for "DNA methylation marks associated with body composition in children from India and the Gambia - findings from the EMPHASIS study"

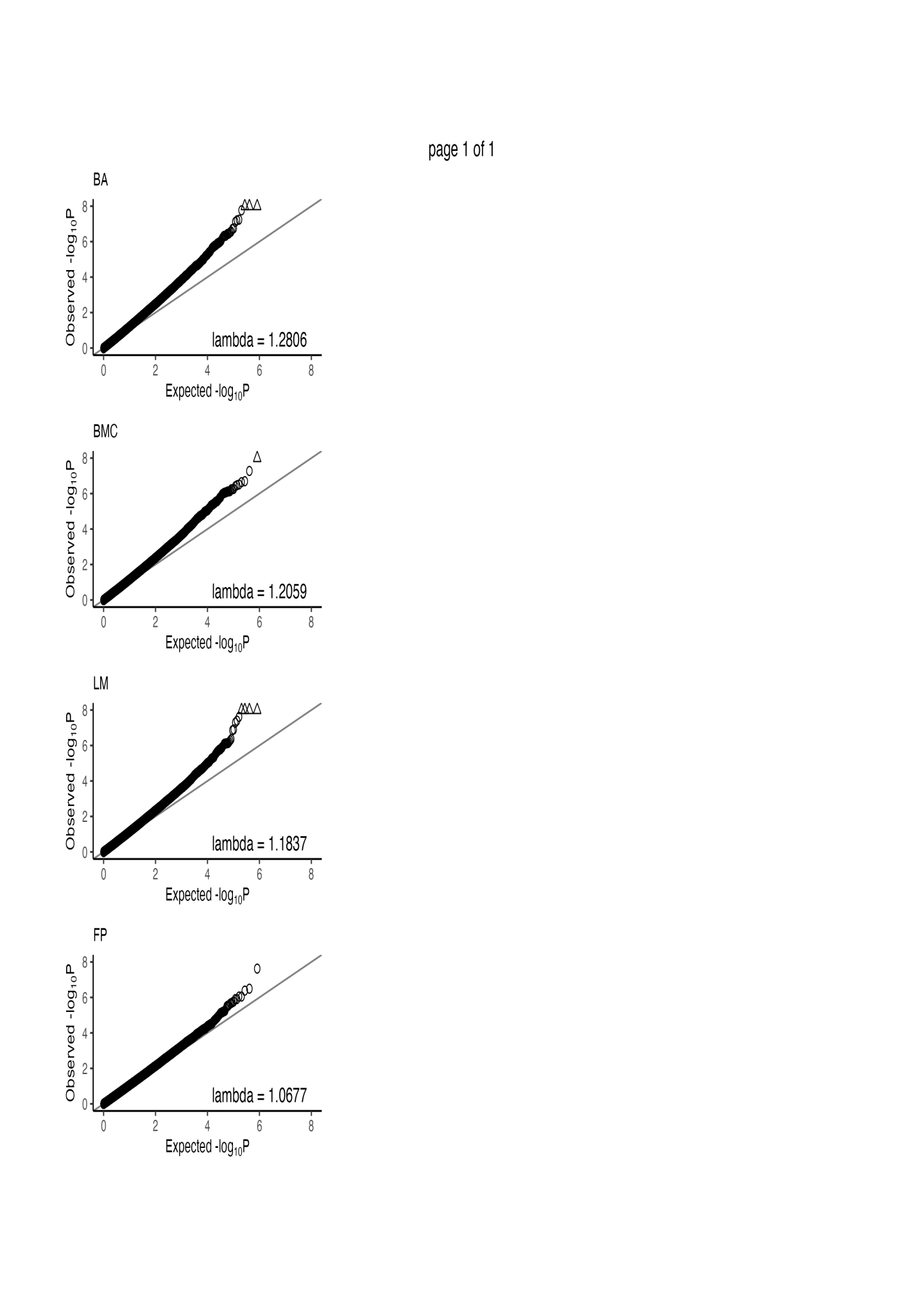

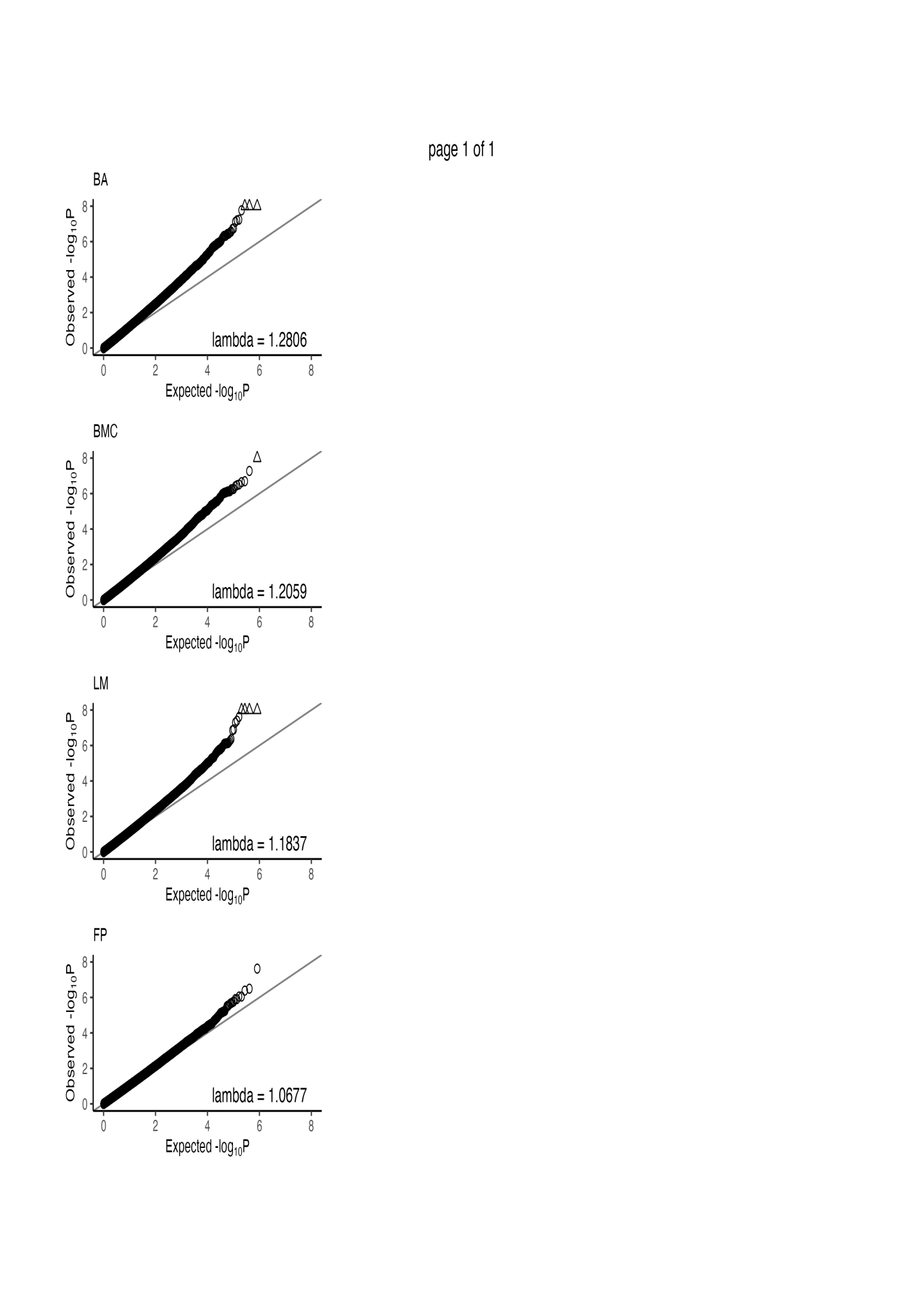

**Supplementary Figure 1:** Quantile-Quantile (Q-Q) plots of EWAS analyses with genome-wide significant associations in the Indian cohort. Lambda indicates the genomic inflation factor. Triangles indicate data points outside the plot area.

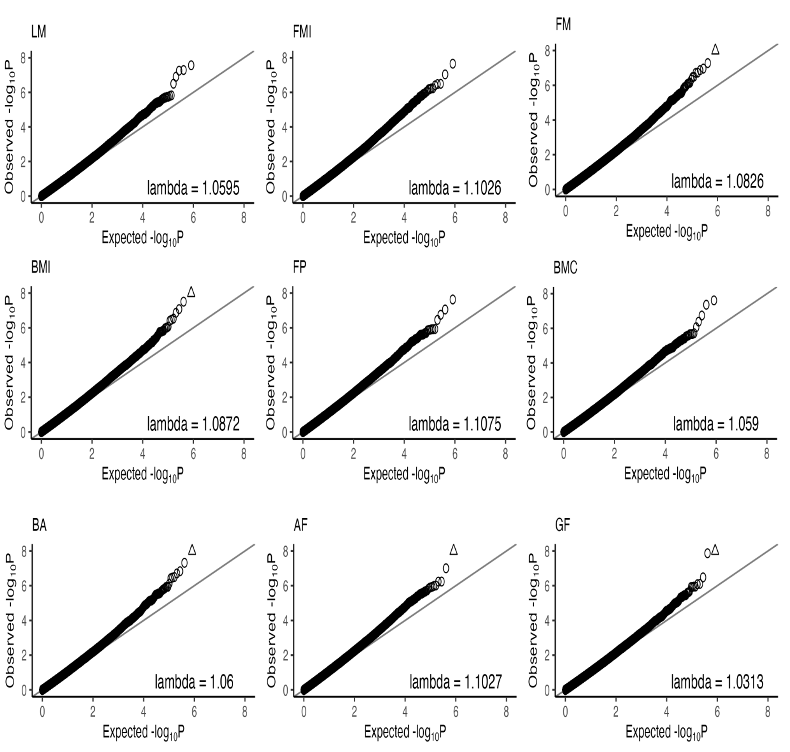

**Supplementary Figure 2:** Quantile-Quantile (Q-Q) plots of EWAS analyses with genome-wide significant associations in the Gambian cohort. Lambda indicates the genomic inflation factor. Triangles indicate data points outside the plot area.
